# Identification of Variable and Joining germline genes and alleles for Rhesus macaque from B-cell receptor repertoires

**DOI:** 10.1101/254599

**Authors:** Wei Zhang, Xinyue Li, Longlong Wang, Jianxiang Deng, Liya Lin, Lei Tian, Jinghua Wu, Huanming Yang, Jian Wang, Ping Qiu, Tong-Ming Fu, Nitin K. Saksena, I-Ming Wang, Xiao Liu

## Abstract

The Rhesus macaque is a valuable preclinical animal model to estimate vaccine effectiveness, and is also important for understanding antibody maturation and B-cell repertoire evolution responding to vaccination; however, incomplete mapping of rhesus immunoglobulin germline genes hinders the research efforts. To address this deficiency, we sequenced B-cell receptor (BCR) repertoires of 75 India Rhesus macaques. Using a bioinformatic method that has been validated with BCR repertoire analysis of three human donors, we were able to infer rhesus Variable (V) and Joint(J) germline alleles, identifying a total of 122 V and 20 J germline alleles. Importantly, 91 V and 13 J alleles were novel, and 40 V and 13 J genes were found at a novel genome region that has not been previously recorded. The novelty of these newly identified alleles was supported by two observations. Firstly, 50 V and 5 J novel alleles were observed in whole genome sequencing data of 10 Rhesus macaques. Secondly, using alignment reference including the novel alleles, the mutation rate of rearranged repertoires was significant declined in 9 other irrelevant samples, and all our identified novel V and J alleles were 100% identity mapped by rearranged repertoire data. These newly identified novel alleles, along with previous reported alleles, provide an important reference for future investigations of rhesus immune repertoire evolution, in response to vaccination or infection. In addition, the method outlined in our study offered an example to future efforts in identifying novel immunoglobulin alleles.

## Introduction

Due to genomic similarity with humans(1), Rhesus macaques has been widely used as an animal model for various immunological and clinical studies, including human immunodeficiency virus 1 (HIV-1), Epstein-Barr virus (EBV), cytomegalovirus (CMV), influenza virus (2-7). It is now feasible for characterizing on a large-scale the immunoglobulin (Ig) repertoires attributed to dramatic improvements in high-throughput sequencing technology. Immune repertoire sequencing (Rep-seq) is a rapidly growing area, and is now commonly being used with applications in studying T cell receptor (TCR)and B cell receptor/antibody (BCR/Ab) repertoires, and for investigating immune responses, prognosis following transplantation, monitoring vaccination responses, identification of neutralizing antibodies and inferring B cell trafficking patterns (8-16). Recently published studies have shown the utility of reopertoire sequencing in the detection of the antibodies (Abs) by Rep-seq technology in Rhesus macaques; in studying vaccine-elicited B cell responses against the HIV primary receptor binding site (3); and in screening of HIV broadly neutralizing antibodies (bNAbs) for HIV-1–infected individuals using bioinformatic methods(17, 18). Antibody production emanates from the rearrangement from multiple sets of similar genes. In this context, it is already known that during the B cell development, both V and J gene segments (light chain) and the V, Diversity(D), and J gene segments (heavy chain) come together to form a functional VL- or VH-region coding sequence through site-specific recombination, also called V(D)J rearrangement. Therefore, we have characterized germline V and J gene sequences, which are critical for analyzing and interpreting Rep-seq data and may serve as a reference database.

Currently, only 23 VH (variable heavy-chain) germline alleles for Rhesus macaques are listed in IMGT database (http://www.imgt.org/). In addition, a study from the Karolinska Institutet, Sweden, extracted 61 genes from the assembled Indian Rhesus genome (termed “KI_genome” hereafter) (3), and including 18 VH alleles from IgM Rep-seq data (termed “KI_IgM”) using bioinformatic methods (19). However, due to the high sequence homology among VH genes, precise and accurate assembly of genome region of VH gene sets are difficult to achieve, especially with insufficient data from alternative source to support the assembly. Moreover, the total number of reported Rhesus VH alleles is much lower than those in human where more than 400 have been reported in IMGT. Thus, it is reasonable to assume that many of VH gene alleles in rhesus macaques have not been identified. Incomplete reference of Rhesus VH and JH (joint heavy-chain) germline alleles resulted in inaccurate assessment on mutational analysis for antibodies, affecting analysis of BCR affinity maturation of clone lineages (20).

To address this insufficiency in rhesus immune gene database, we sequenced IgH repertoires for a total of 75 rhesus monkey samples. To support identification of novel V and J germline alleles, we developed a software IMPre to infer T-cell receptor(TCR) and BCR germline alleles from Rep-seq data (21). By analyzing VH and JH germline alleles for each sample, we were able to acquire the alleles presented in multiple samples. The exercise yielded 122 IgH V and 20 J germline alleles from these samples, including 91 novel VH and 13 JH alleles. These newly identified alleles were validated in two independent datasets, one is whole genome sequencing data from 10 rhesus monkeys and the other is the IgH Rep-seq data of 9 rhesus samples from other projects. We collected all previously reported dataset along with our newly identified rhesus germline genes and alleles as a reference for future immune repertoire research.

## Materials and Methods

### Sample Collection and Data Production

Fifteen Indian Rhesus macaques were collected, and we adhered to the guidelines for the care and use of animals for scientific purposes established by the Singapore National Advisory Committee for Laboratory Animal Research (NACLAR) in November 2004. For each monkey, five time points (one pre-vaccine and four post- dengue vaccine) of peripheral blood (PB) samples were collected, and then peripheral blood mononuclear cells (PBMC) were isolated from the peripheral blood samples. Total RNA was extracted using TRIzol (Invitrogen, Carlsbad, CA, USA), and a Rapid Amplification of cDNA Ends (5’RACE) kit (v2.0, Invitrogen) with primers at the constant region (Supplementary Table S1) were used to amplify IgH repertoire. The experimental details were described in our previously published paper(22). In addition, a restriction Enzyme site was added in the 5’ site of the cDNA, and the Abridged Anchor Primer (AAP) introduced during RACE was cut off after the PCR. Illumina sequencing adaptors were ligated directly to the PCR amplicons without shearing, and the libraries were sequenced on Illumina Miseq platform to produce paired-end (PE) 300 reads

To evaluate the accuracy of method, three human RNA samples were collected and were subjected to multiple PCR with BIOMED-2 primers(23) at the FR1 and C regions to amplify the IgH repertoire. To validate our identified novel germline V and J genes and alleles, whole genome sequencing (WGS) of 10 India Rhesus macaques and IgH Rep-seq (RNA) of another 9 India Rhesus macaques were involved. The raw sequencing data of the former were randomly selected from a published study(10) and downloaded from NCBI Sequence Read Archive (SRA; https://www.ncbi.nlm.nih.gov/sra). The data of the latter include two sources, 4 samples were from other published studies(4, 19), and the other 5 samples were from our another HIV vaccine project which were also amplified by 5’RACE kit (v2.0, Invitrogen) and sequenced by Illumina Hiseq 2000 platform with PE150 reads.

For human samples used in this this study were prospectively reviewed and approved by a duly constituted ethics committee (the institutional review board on bioethics and biosafety of BGI ethical approval).

### VH and JH Germline Allele Inference for Each Sample

Firstly, raw sequencing data was processed and the paired-end reads were merged using IMonitor(13). According to the IgH leader sequences of Rhesus in the IMGT database, full or partial leader sequences observed in the merged sequences were trimmed. As for each sample, we used our previously developed software IMPre to predict candidate IgH V and J germline alleles from the rearranged IgH repertoires(21). IMPre is based on the V/D/J rearrangement mechanism to infer original germline gene segment, consisting of four steps, including data processing, clustering, assembly, and optimization. It is rigorous in eliminating the false positive sequences. With this, dozens of candidate germline alleles were generated for each sample.

### VH and JH Germline Processing

Firstly, to ensure the exact sequence length inferred for V and J germline alleles, we modified the terminal ends of sequences by adding or trimming few nucleotides. Moreover, the inferred germline alleles were mapped to human and previously reported Rhesus germline alleles, and the nearest human or Rhesus germline alleles were identified for each candidate allele. Firstly, for comparing with the nearest allele, we deleted the extra nucleotides of candidate allele at the terminal ends, or added the most frequent sub-sequence in the sample’s raw data if the length of inferred allele was less than that of the nearest one. And, secondly, we combined the inferred V or J germline alleles that came from one individual. Those alleles were clustered by sequence similarity by QTClust algorithm(24). The allele present in most samples was as defined as the cluster center, and other allele was classified and integrated to this cluster if the mismatch was less than 3bp. The alleles from one cluster were regarded from one gene segment, and at least two alleles were truly correct. The allele observed in most samples was regarded as major allele, whereas the less dominating alleles were regarded as minor. Thus for each cluster only major and minor alleles were retained, and others were filtered out. Thirdly, the allele that did not contain conservative motif was filtered out. The conservative motif for VH germline allele was the conservative amino acid “YYC” during the start of CDR3, and the motif for JH germline allele was “FGXG” (‘X’ could be any amino acid) near the end of CDR3 region. Fourthly, the allele presented across multiple samples were considered as final germline allele. As for the V gene, we only retained the allele observed in more than four samples, for the VH4 allele retention was based on if observed in more than two individuals, and for the J gene, retention of allele was based on its presence in more than two individuals also.

### Nomenclature for Novel Germline Genes and Alleles

We named these sequences using previously reported KI_genome alleles(3) and the IMGT collaboration nomenclature rules. As for VH germline genes and alleles, we used the nomenclature rules of the KI_genome, whereas the nomenclature rules of the IMGT were used in classifying the JH alleles. We also added a letter and a number that was used to distinguish the final annotations when a complete genome assembly became available for future use, with annotation such as “VH1-1A*01”. Our inferred VH and JH alleles were aligned to IMGT, KI-genome and Human germline alleles, and we selected the nearest mapped allele for each inferred allele. Besides, our inferred novel VH alleles were aligned to the Macaca mulatta (Rhesus monkey) genome (ftp://ftp.ncbi.nih.gov/genomes/Macaca_mulatta). We used five steps to nomenclate the sequences, as discussed below.

**1.** The sequence’s gene family were identified according to the nearest allele (e.g., VH1, IGHJ1).

**2.** In our previous study, we analyzed the allele differences within and between genes(21), and found the differences for most alleles within gene were less than 7 mismatches for the VH and 5 mismatches for the JH. Thus, if mismatches between the inferred and nearest allele were less than 7bp (for J: <=5bp), the inferred allele was classified into the gene of the nearest allele. The novel allele with fewer mismatches was prior to offer a smaller number in the name (e.g., VH1.1A*02 (1 mis-match), and VH1.1A*03 (2 mismatches)).

**3.** If the mismatches between the inferred and nearest allele were more than 7bp (for J: >5bp), we then compared the genome position that inferred allele mapped with at the very position previous alleles mapped to. If the mapped location in the Rhesus genome was different from that of the previous alleles, inferred allele was defined as a novel gene, and we nomenclated as “gene family-new number A*01“, where ‘new-number’ is a number (e.g., VH2.70A*01).

**4.** Subsequently, if the mismatches between the inferred and nearest allele was more than 7bp (for J: >5bp), and the mapped location at Rhesus genome was same as that of one previous allele, the inferred allele was defined as a novel gene, we subsequently named them as “gene family-number B*01“, where ‘number’ was same as the gene of nearest allele. If we generated another novel gene, the letter ‘B’ was changed to ‘C’, and so on. However, few nearest alleles originated from Human, in this case, the alleles were named as per the rules described in step 4.

**5.** Finally, our inferred novel alleles and all previously reported germline alleles were collected together. All alleles added a lowercase to represent the origin. ‘a’: from IMGT, ‘b’: from KI_genome, ‘c’: from our inferred novel alleles, ‘d’: from KI_IgM.

### Validation by Rhesus genome data

Ten whole-genome sequencing (WGS) India Rhesus data(10) were downloaded from NCBI Sequence Read Archive (SRA). The raw data were converted into fastq paired-end reads using sratoolkit(v.2.8.2-1) and then they were aligned against the genome reference of Indian Rhesus macaque using the software BWA. The reads mapped to chromosome 7, chromosome 7 un-localized scaffolds, unplaced genomic scaffolds and unmapped reads were extracted for subsequent analysis.

To explore whether our inferred VH germline alleles existed in the WGS data, those extracted reads were aligned to our inferred VH germline alleles using the software BWA. The reads with more than 1 mismatch in comparison to the reference sequence was filtered out, and the coverage for each VH allele was calculated. The resulting sequence alignments were viewed by the Integrative Genomics Viewer (IGV, v2.3.97)(25). To explore whether our inferred JH germline alleles existed in the WGS data, the extracted reads were used to search our inferred JH germline alleles. The reads included full-length JH allele sequence and no mismatch was allowed. The number of eligible reads for each sample was recorded.

### Validation by other IgH Rep-seq data

For this, we downloaded the previously published IgH raw Rep-seq data on four rhesus samples(4, 19), along with collecting five vaccined samples from our another project. All IgH repertoire data were processed and analyzed with the tool IMonitor(13). Rearranged sequences were mapped to the Rhesus V and J germline gene segments and the mismatches between them were regarded as mutations. The parameters of IMonitor used were: -vif 70 -jif 70. The sequence mapped to both V and J germline alleles was retained for further analysis. We constructed two reference datasets used in the alignment step. One reference dataset comprised of all of the previously reported VH and JH germline alleles, whereas the second reference dataset consisted of our novel and previously described alleles. In comparison, raw data of each sample were aligned to both reference datasets for further analysis. To reduce the effect of sequencing error, the sequence present in the sample only once was filtered. Finally, we selected the same sequences mapped to both reference datasets and were considered for mutation calculation. The mutation rate was defined as the total mismatch number divided by total alignment nucleotides amount.

### Statistics analysis

In this study, Mann-Whitney two-tailed test (by Wilcoxon test in R) was used to test and calculate the p value. P values were corrected by False Discovery Rate (FDR) for multiple tests.

## Results

### Systematic design and Evaluation

As shown in Fig. 1A, we utilized our newly developed tool IMPre to infer and analyzing novel germline genes and alleles for each sample from the Rep-seq data(21). IMPre includes data processing, clustering, assembling and optimization steps to get the candidate novel germline alleles. Following the processing of 75 rhesus samples using the IMPre, the terminal nucleotides of inferred alleles were modified in terms of human and previously described Rhesus germline alleles. After that, all the alleles from one individual were combined together and clustered by sequence similarity. Only two alleles, that were used by most samples, were retained. The inferred alleles that comprised the VH or JH conservative amino acid motif, and that presented in multiple samples were regarded as the final germline alleles. Finally, the novel genes and alleles were named according to the KI-genome and IMGT collaboration nomenclature rules by comparing the previously reported Rhesus and human germline alleles, and the Rhesus genome (see METHODS section for more details).

**Figure 1.**
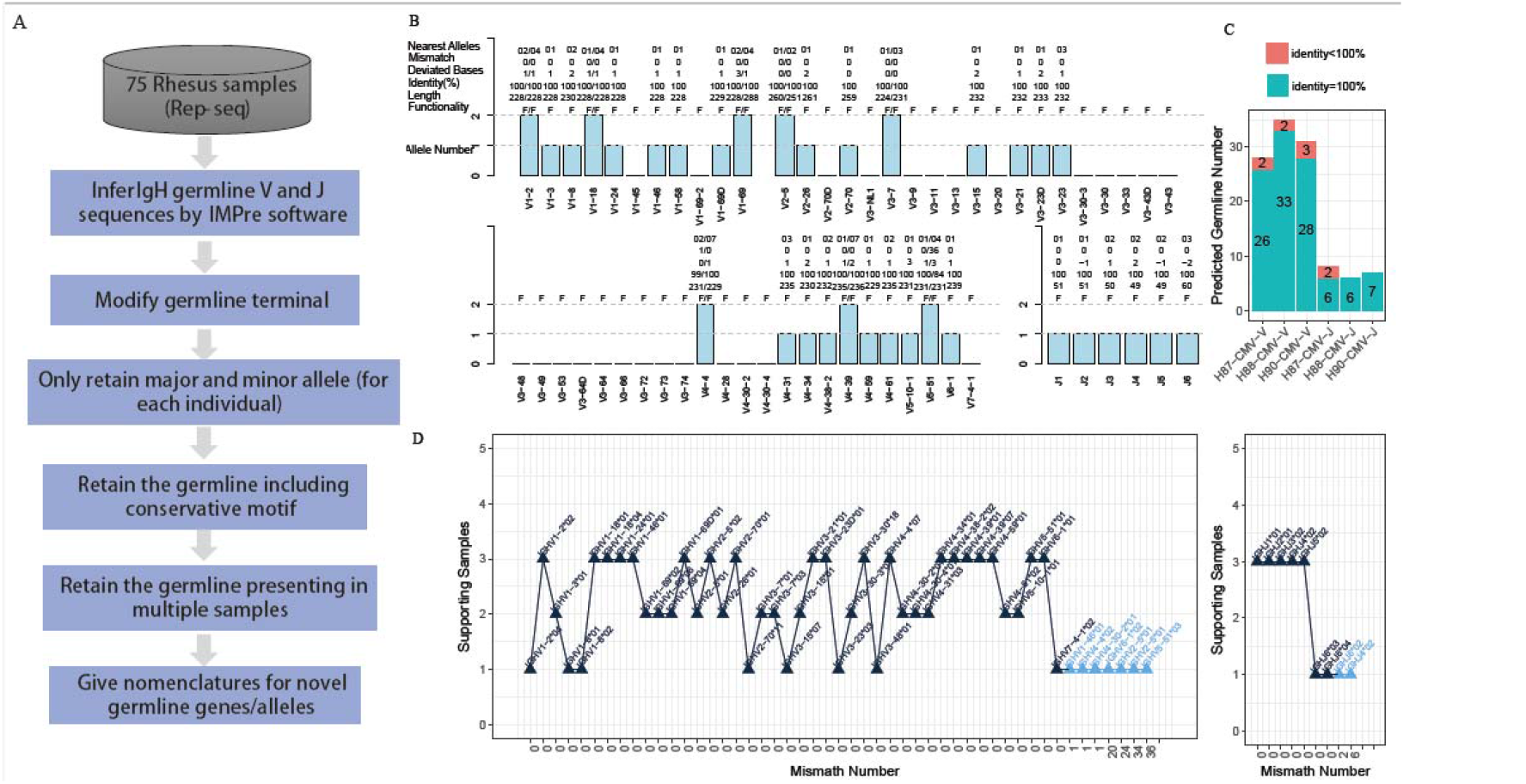
Schema of the overall design of the method and detailed evaluation of predicted alleles for three human IgH samples. (A). Overall pipeline of inferring V and J germline alleles from rearranged IgH repertoires. (B). Inferred alleles for H88-LS sample. The predicted alleles by IMPre were aligned to human IgH germline alleles, and the accuracy was evaluated. The V3 alignments only accounted for 3.7% of the raw sample sequences; therefore, most V3 genes did not have any allele prediction. (C). Identity between predicted alleles and human IgH alleles. (D). Relationship between mismatch number and supporting samples for inferred alleles from three samples. Supporting sample- the number of samples observed the inferred allele.

To evaluate the accuracy of our strategy that infers germline alleles from multiple samples, we applied it to the 3 human IgH samples because human germline genes are pretty complete so far. The VH and JH germline alleles of three samples inferred by IMPre were showed in Fig. 1B and Fig. S1. Almost all of the inferred VH and JH alleles were completely consistent with the true alleles, with the exception of some inferred alleles showing <100% identity. (Fig. 1C), implying the robust performance of IMPre. Inferred germline alleles from the three known samples were combined and processed by our strategy. Consequently, forty-nine VH and nine JH inferred germline alleles were obtained and shown in Fig. 1D. We observed that 34 of 49 VH alleles were supported by at least two individuals, and all of them perfectly matched with the human germline alleles. Six of nine J alleles were supported by at least two individuals, and they were all consistent with human alleles. Together, these results show excellent concordance in results, along with the capacity of our method in referring and inferring germline alleles from multiple individuals.

### Prediction of Rhesus Monkey Novel Germline J Alleles

We designed primers in the conservative C region to capture the IgH rearranged repertoires (of 75 India-Rhesus samples) using high-throughput sequencing (Miseq, paired-end 300bp). We first utilized the IMPre to infer VH and JH germline alleles and then combined them with further processing (see the details in the Methods section). Finally, we inferred 20 JH germline alleles from a total of 75 Indian-Rhesus samples, of which 7 of them were the same as the Rhesus JH germline alleles in IMGT database (Fig. 2A, Table 1). There were 8 JH alleles in IMGT database, with the exception of the allele IGHJ5-2*01 which was missing in our analysis. The high consistency between our results and the IMGT database further validate that our method for inferring germline alleles performs robustly and accurately, resulting in high quality of the inferred alleles.

**Figure. 2:**
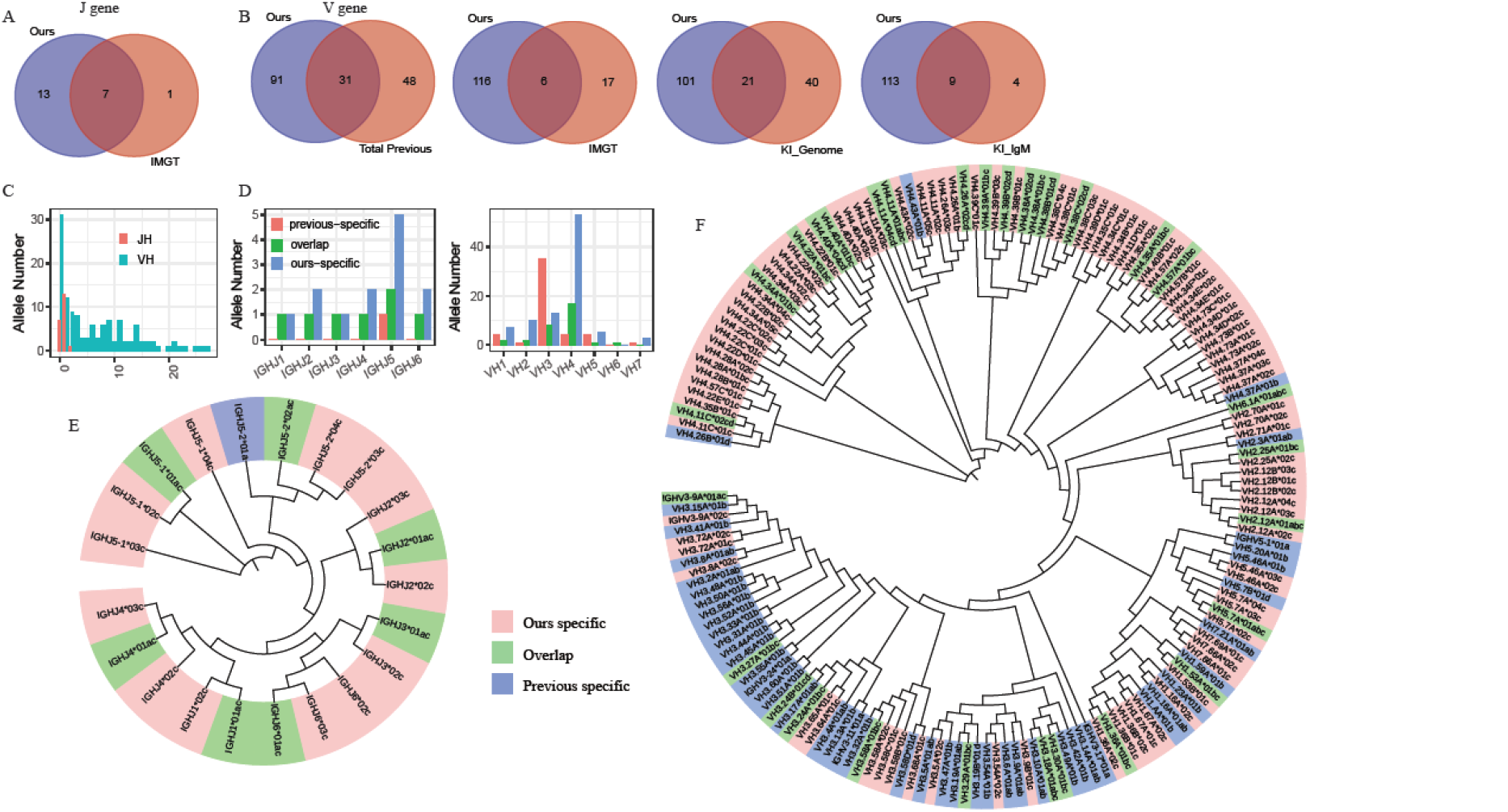
Predicted Rhesus monkey germline genes and alleles. (A). Venn diagram shows the comparison of allele number between our identified and IMGT JH alleles. (B). Venn diagram shows the comparison of allele number between our identified and previous reported VH alleles. The previous reported alleles include three sources: IMGT, KI_genome, KI_IgM. (C). The distribution of mismatch number between our identified alleles and the mapped nearest reported alleles. (D). VH and JH germline allele number were distributed among all the gene family. Previous specific: present only in previously reported alleles; overlap: present in both reported and our identified alleles; our specific (novel): only present in our identified alleles. (E). An unrooted circular phylogram showing the clustering relationships between previously reported and newly identified JH alleles. The lowercase letter at the allele’s terminus indicates the origin of the sequence, a: IMGT; b: KI_genome; c: our identified; d: KI_IgM. (F). An unrooted circular phylogram for VH alleles. The VH and JH alleles’ sequences were processed by MUSCL(http://www.ebi.ac.uk/Tools/msa/muscle/) and the tree was visualized by the Interactive Tree of Life50 (version 3.6, http://itol.embl.de/).

**Table 1.**
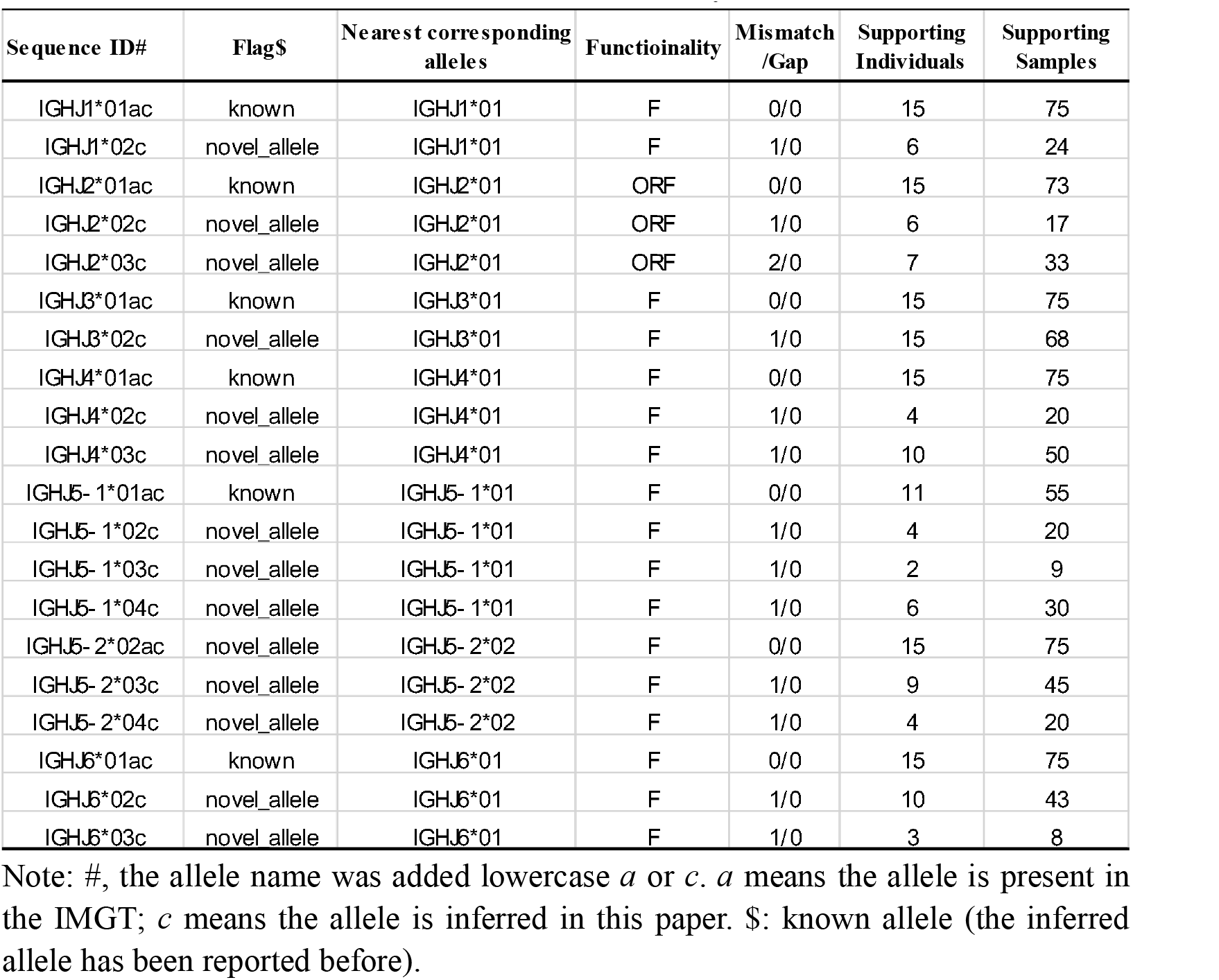
Identified novel JH alleles in the Rhesus monkey

Apart from the 7 overlapping alleles with IMGT, the other 13 novel alleles encompass the full IgH J genes from JH1 to JH6 (Fig. 2D). There was only one nucleotide difference between the novel alleles and their nearest previous alleles, except that the allele IGHJ2*03c showed two mismatches with the allele IGHJ2*01 (Fig. 2C). The 7 alleles that also existed in the IMGT were observed in all 15 individuals, probably because those alleles are commonly used in the population. All novel alleles were present in more than two individuals and in more than 8 samples, and four of the novel alleles were observed in more than 10 individuals (Table 1). Finally, we named all our novel inferred alleles using the IMGT collaboration nomenclature rules. We combined the novel alleles and the alleles in the IMGT database and displayed them in a phylogenetic tree on the basis of sequence homology (Fig. 2E). We also provided all JH alleles’ sequences by FASTA format in the supplementary information.

### Prediction of Rhesus Monkey Novel Germline V Genes and Alleles

Up to now, there are three reported datasets of VH germline alleles: 23 alleles in IMGT database, 61 alleles extracted from Rhesus genome(3), and 13 alleles (only alleles presented in at least 2 monkeys) identified from IgM repertoires using bioinformatic method(19). From 75 Indian Rhesus samples we inferred a total of 122 VH germline alleles, of which 91 were novel (Table 2, 3 and Table S2). The remaining 31 inferred alleles were completely consistent with one or more of the three reported datasets (Table S2), including 6, 21 and 9 alleles overlapped with the IMGT database, KI_genome, and KI_IgM, respectively (Fig. 2B). The 31 reported alleles were distributed across all VH gene families (2 in VH1, 2 in VH2, 8 in VH3, 17 in VH4, 1 in VH5 and 1 in VH6), with the exception of VH7. Likewise, the 91 novel alleles encompassed all VH genes (including 7 in VH1, 10 in VH2, 13 in VH3, 53 in VH4, 5 in VH5 and 3 in VH7), with the exception of VH6 (Fig. 2D). The number of inferred VH4 alleles was the most abundant amongst all VH genes, which could be attributed to the large number of VH4-origin sequences from the raw Rep-seq data.

**Table 2.**
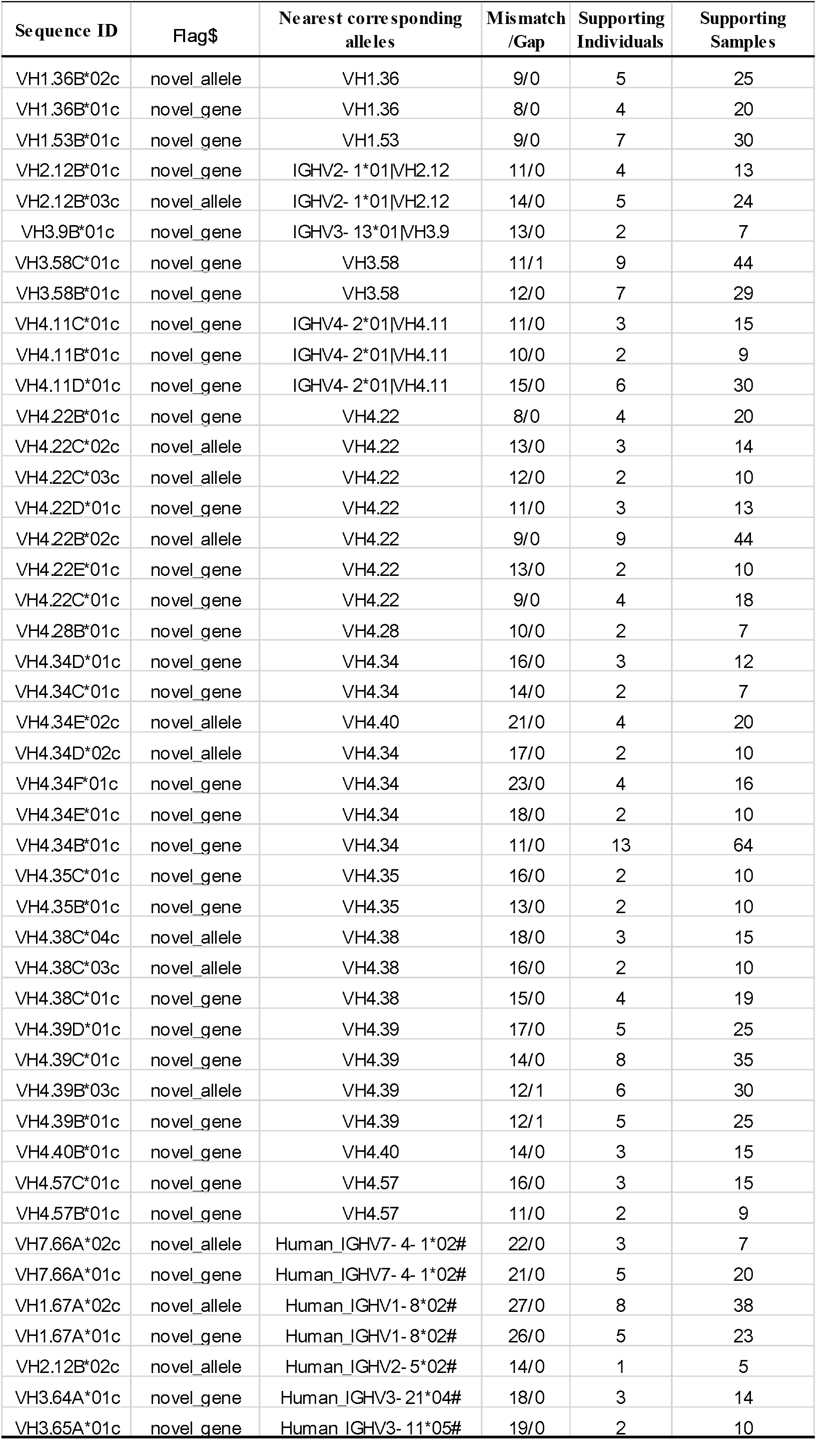
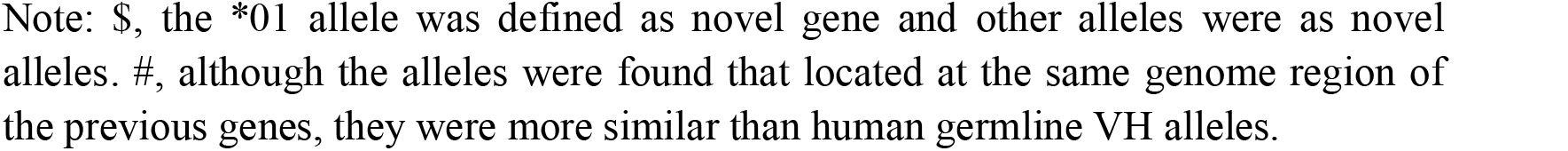
Predicted novel V genes and alleles in the Rhesus monkey (mismatch >7bp, and located at the same genome region of the previous genes)

| Sequence ID | Flag\$ | Nearest corresponding alleles | Mismatch /Gap | Supporting Individuals | Supporting Samples |
| --- | --- | --- | --- | --- | --- |
| VH1.36B*02c | novel allele | VH1.36 | 9/0 | 5 | 25 |
| VH1.36B*01c | novel gene | VH1.36 | 8/0 | 4 | 20 |
| VH1.53B*01c | novel gene | VH1.53 | 9/0 | 7 | 30 |
| VH2.12B*01c | novel gene | IGHV2- 1*01\|VH2.12 | 11/0 | 4 | 13 |
| VH2.12B*03c | novel allele | IGHV2- 1*01\|VH2.12 | 14/0 | 5 | 24 |
| VH3.9B*01c | novel gene | IGHV3- 13*01\|VH3.9 | 13/0 | 2 | 7 |
| VH3.58C*01c | novel gene | VH3.58 | 11/1 | 9 | 44 |
| VH3.58B*01c | novel gene | VH3.58 | 12/0 | 7 | 29 |
| VH4.11C*01c | novel gene | IGHV4- 2*01\|VH4.11 | 11/0 | 3 | 15 |
| VH4.11B*01c | novel gene | IGHV4- 2*01\|VH4.11 | 10/0 | 2 | 9 |
| VH4.11D*01c | novel gene | IGHV4- 2*01\|VH4.11 | 15/0 | 6 | 30 |
| VH4.22B*01c | novel gene | VH4.22 | 8/0 | 4 | 20 |
| VH4.22C*02c | novel allele | VH4.22 | 13/0 | 3 | 14 |
| VH4.22C*03c | novel allele | VH4.22 | 12/0 | 2 | 10 |
| VH4.22D*01c | novel gene | VH4.22 | 11/0 | 3 | 13 |
| VH4.22B*02c | novel allele | VH4.22 | 9/0 | 9 | 44 |
| VH4.22E*01c | novel gene | VH4.22 | 13/0 | 2 | 10 |
| VH4.22C*01c | novel gene | VH4.22 | 9/0 | 4 | 18 |
| VH4.28B*01c | novel gene | VH4.28 | 10/0 | 2 | 7 |
| VH4.34D*01c | novel gene | VH4.34 | 16/0 | 3 | 12 |
| VH4.34C*01c | novel gene | VH4.34 | 14/0 | 2 | 7 |
| VH4.34E*02c | novel allele | VH4.40 | 21/0 | 4 | 20 |
| VH4.34D*02c | novel allele | VH4.34 | 17/0 | 2 | 10 |
| VH4.34F*01c | novel gene | VH4.34 | 23/0 | 4 | 16 |
| VH4.34E*01c | novel gene | VH4.34 | 18/0 | 2 | 10 |
| VH4.34B*01c | novel gene | VH4.34 | 11/0 | 13 | 64 |
| VH4.35C*01c | novel gene | VH4.35 | 16/0 | 2 | 10 |
| VH4.35B*01c | novel gene | VH4.35 | 13/0 | 2 | 10 |
| VH4.38C*04c | novel allele | VH4.38 | 18/0 | 3 | 15 |
| VH4.38C*03c | novel allele | VH4.38 | 16/0 | 2 | 10 |
| VH4.38C*01c | novel gene | VH4.38 | 15/0 | 4 | 19 |
| VH4.39D*01c | novel gene | VH4.39 | 17/0 | 5 | 25 |
| VH4.39C*01c | novel gene | VH4.39 | 14/0 | 8 | 35 |
| VH4.39B*03c | novel allele | VH4.39 | 12/1 | 6 | 30 |
| VH4.39B*01c | novel gene | VH4.39 | 12/1 | 5 | 25 |
| VH4.40B*01c | novel gene | VH4.40 | 14/0 | 3 | 15 |
| VH4.57C*01c | novel gene | VH4.57 | 16/0 | 3 | 15 |
| VH4.57B*01c | novel gene | VH4.57 | 11/0 | 2 | 9 |
| VH7.66A*02c | novel allele | Human_IGHV7- 4- 1*02# | 22/0 | 3 | 7 |
| VH7.66A*01c | novel gene | Human_IGHV7- 4- 1*02# | 21/0 | 5 | 20 |
| VH1.67A*02c | novel allele | Human_IGHV1- 8*02# | 27/0 | 8 | 38 |
| VH1.67A*01c | novel gene | Human_IGHV1- 8*02# | 26/0 | 5 | 23 |
| VH2.12B*02c | novel allele | Human_IGHV2- 5*02# | 14/0 | 1 | 5 |
| VH3.64A*01c | novel gene | Human_IGHV3- 21*04# | 18/0 | 3 | 14 |
| VH3.65A*01c | novel gene | Human_IGHV3- 11*05# | 19/0 | 2 | 10 |

We named novel alleles according to the rules consistent with the KI_genome, because there were more VH genes in the KI_genome than in the IMGT. Our inferred alleles were mapped to the IMGT, KI_genome and human germline alleles. If the mismatch number between inferred allele and its nearest allele was more than 7bp that was found the difference boundary for most VH genes(21), the inferred allele was defined as a novel gene. We found 40 novel genes consisting of 56 alleles (Table 2 and 3). These novel genes were also mapped to the reference of Indian Rhesus genome, and 8 of them were located in the novel regions of the chromosome compared with the previous germline gene (Table 3). One novel gene VH3.68A was located on chromosome 7, two genes were located on the un-localized scaffolds of chromosome 7, and other five genes were located at unplaced genomic scaffolds. Interestingly, among the novel predictions, 9 novel alleles were mapped to human VH germline alleles (Table 2 and 3), which implies that previously reported Indian Rhesus VH genes are incomplete. We also observed several novel alleles bearing a gap with the nearest alleles from the KI_genome. Finally, we combined our novel alleles and all of the previously reported alleles to generate a phylogenetic tree on the basis of their conserved sequences (Fig. 2F). We provide all of those VH alleles’ sequences in FASTA format in Supplementary data.

**Table 3.**
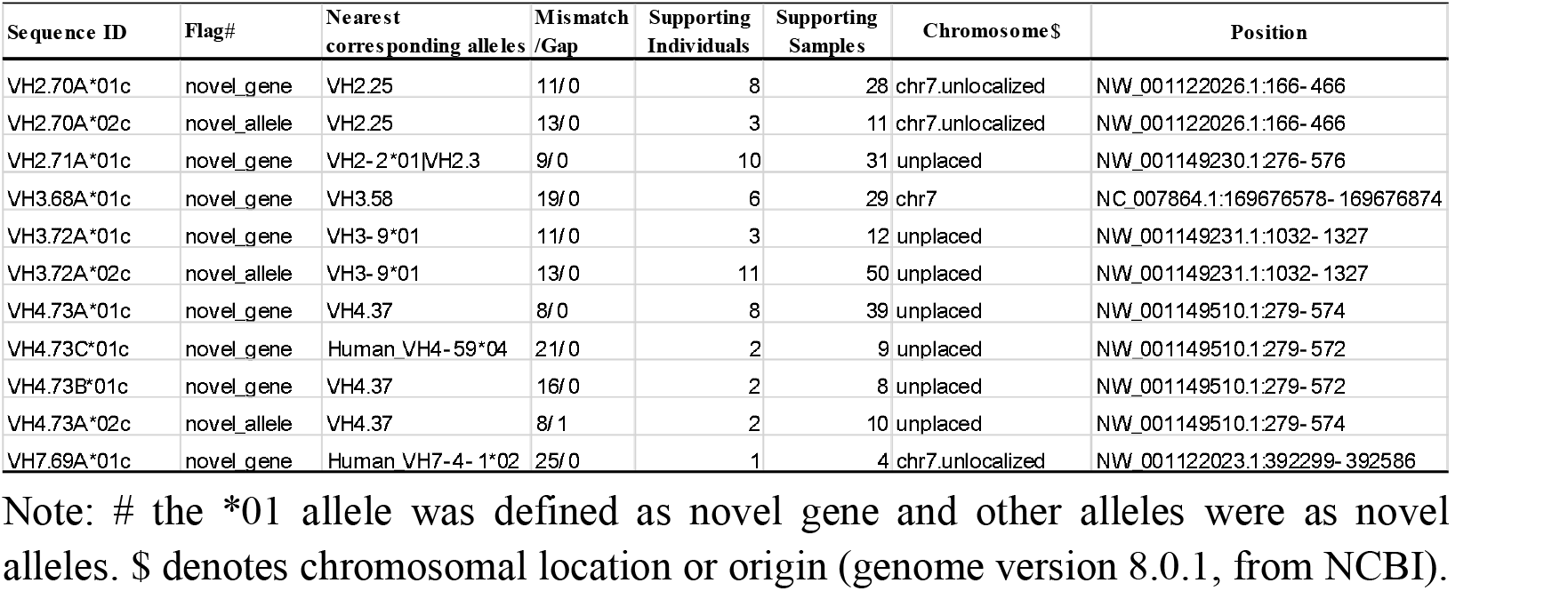
Predicted novel V genes and alleles in the Rhesus monkey (Mismatch>7bp, and located at the novel genome region)

### Validation of novel genes and alleles by 10 India Rhesus genome data

To validate the accuracy of our inferred novel germline alleles, we tried to find whether these alleles existed in 10 Indian Rhesus genome data. The Raw reads of the whole genome sequences were firstly aligned against the Rhesus genome reference, and the reads mapped to chromosome 7, chromosome 7 un-localized scaffolds, unplaced genomic scaffolds were extracted. Moreover, the unmapped reads were also extracted to align against the newly identified alleles. Only one mismatch for the VH alleles, and no mismatch for the JH alleles were allowed for the alignment steps and coverage calculation. The VH allele’s coverage being more than 95% or the JH allele which was completely included in a read was regarded as existing in the Rhesus genome. We found that 73 out of 122 (60%) VH alleles were present in at least one Rhesus genome data, and 50 of these 73 alleles were novel germline alleles (Fig. 3A). Likewise, of 20 inferred JH germline alleles, 12 existed in the Rhesus genome data and 5 of them were novel alleles (Fig. 3A). Actually, many alleles were found in multiple Rhesus monkeys and the number of supporting Rhesus monkeys are displayed in Fig. 3B and Fig. S2. Eighteen (35%) VH novel alleles existed in more than four monkeys, while 32(64%) existed in 1 to 4 monkeys (Fig. 3B). One JH novel allele was observed in eight monkeys, whereas four alleles were observed in 1-2 monkeys. As expected, most of our inferred VH and JH alleles that were previously reported presented in multiple monkeys (Fig. S2). To view the details of the coverage and alignment, we utilized the tool Integrative Genomics Viewer (IGV) to display several VH alleles. For examples, VH1.16A*02c was found in 10 Rhesus genome data and displayed 100% coverage in almost all samples (Fig. 3C). We also took Rhesus SRR1927133 for example to show the information of mapped reads. Four other VH alleles’ coverage were displayed in Fig. S3 and S4, and two VH alleles’ alignment reads are shown in Fig. S5. IGHJ5-1*02c was found in 2 Rhesus monkeys, and IGHJ5-1*04c was found in 8 monkeys. Their mapped reads are shown in Fig. 3E. Therefore, the high coverage for novel VH and JH alleles imply that our inferred alleles are reliable.

**Figure 3:**
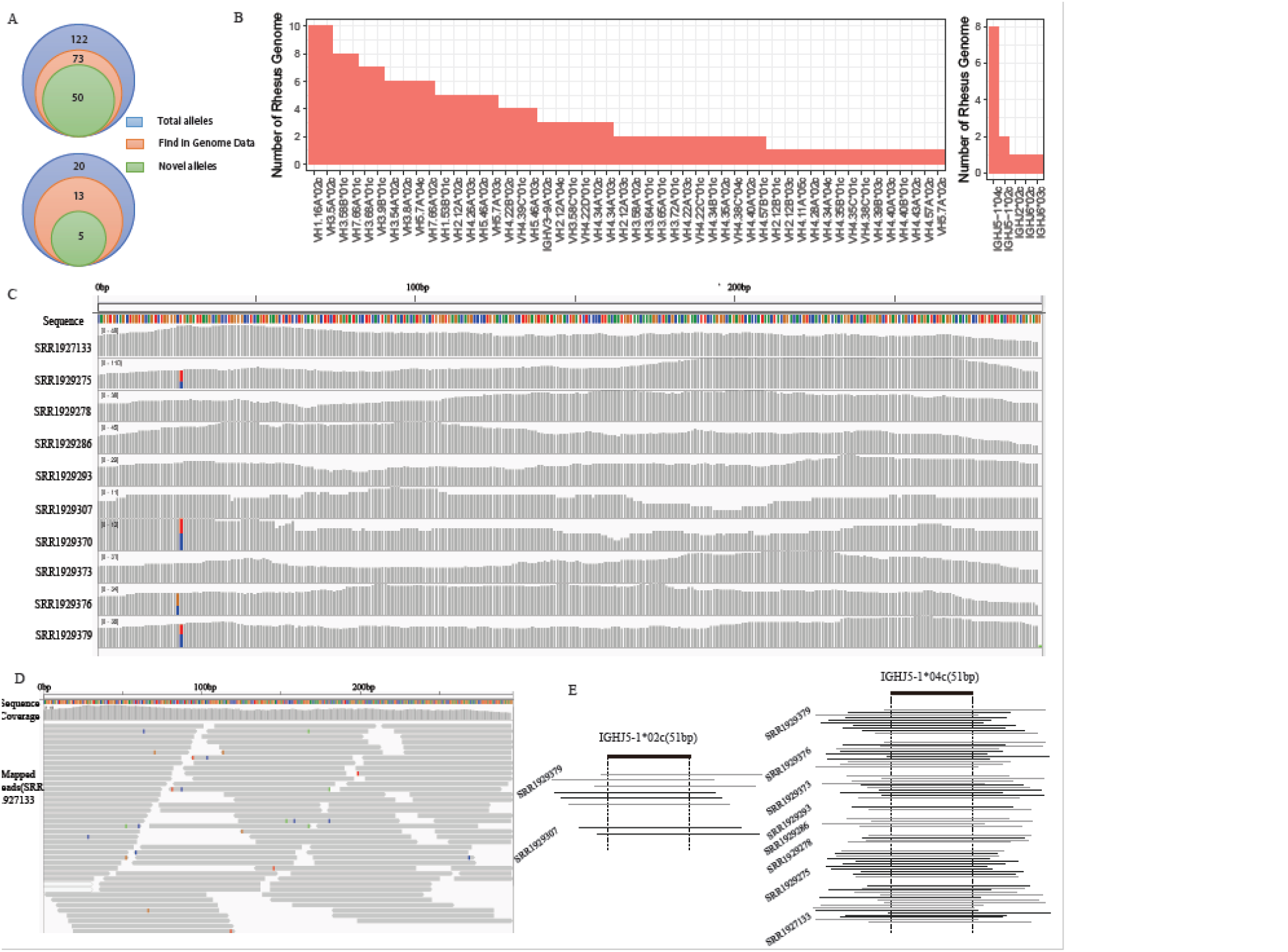
Germline genes and alleles validation in Rhesus genome data. Reads from whole genome sequencing were mapped to our inferred germline alleles. (A). The number of our inferred germline alleles present in the genome data. (B). The number of monkeys which supported the VH or JH novel alleles. (C). Coverage of VH1.16A*02c in 10 Rhesus genome data. This figure was implemented by IGV (v2.3.97). Colors represent different type of bases (A,C,G,T). (D). Visualization of detailed reads alignment for VH1.16A*02c. The reads came from one of the Rhesus SRR1927133. Mismatches between the reads and reference are highlighted in different colors. The figure was drawn in the IGV tool. (E). Visualization of reads covering novel alleles IGHJ5-1 *04c and IGHJ5-1*02c. Black bars represent reads from the plus chain, and the grey bars represent reads from the minus chain.

### Validation of novel genes and alleles by other IgH Rep-seq data

We utilized other IgH Rep-seq data to further evaluate the novel genes and alleles, which included three sources: Christopher S. et al. (F128) (4), Martin M. C. et al., (3 Indian Rhesus)(19), and 5 HIV-vaccine samples from a unpublished project. The rearranged IgH repertoire sequences of the 9 samples were mapped to the two reference datasets: previous reference (V germline alleles from the IMGT, KI_genome, KI_IgM) and total reference that we termed the MacVJ hereafter (previously described + our novel alleles). Comparing with the alignment with previously described reference sequences, the identities between sequences and MacVJ sequences were much higher for both VH and JH genes (Fig. S6A). During the development of BCRs, creating hyper-mutation is one of essential step for improving the affinity maturation to efficiently react with auto- or external antigens, and so an accurate identification of mutation is critical for BCR analyses, such as lineage analysis(20) and active clone determination(26). Thus, more VH and JH germline alleles need be obtained for the mutation identification. To view novel germline alleles’ effect on mutation analysis, we calculated mutation rates of each VH and JH gene families in both previously described sequences and MacVJ sequences, and found the rate significantly declined across all samples and most gene families using MacVJ (Fig. 4A). This improve the detection of mutations in the newly identified alleles, which is vital for the BCR repertoire analysis.

**Figure 4:**
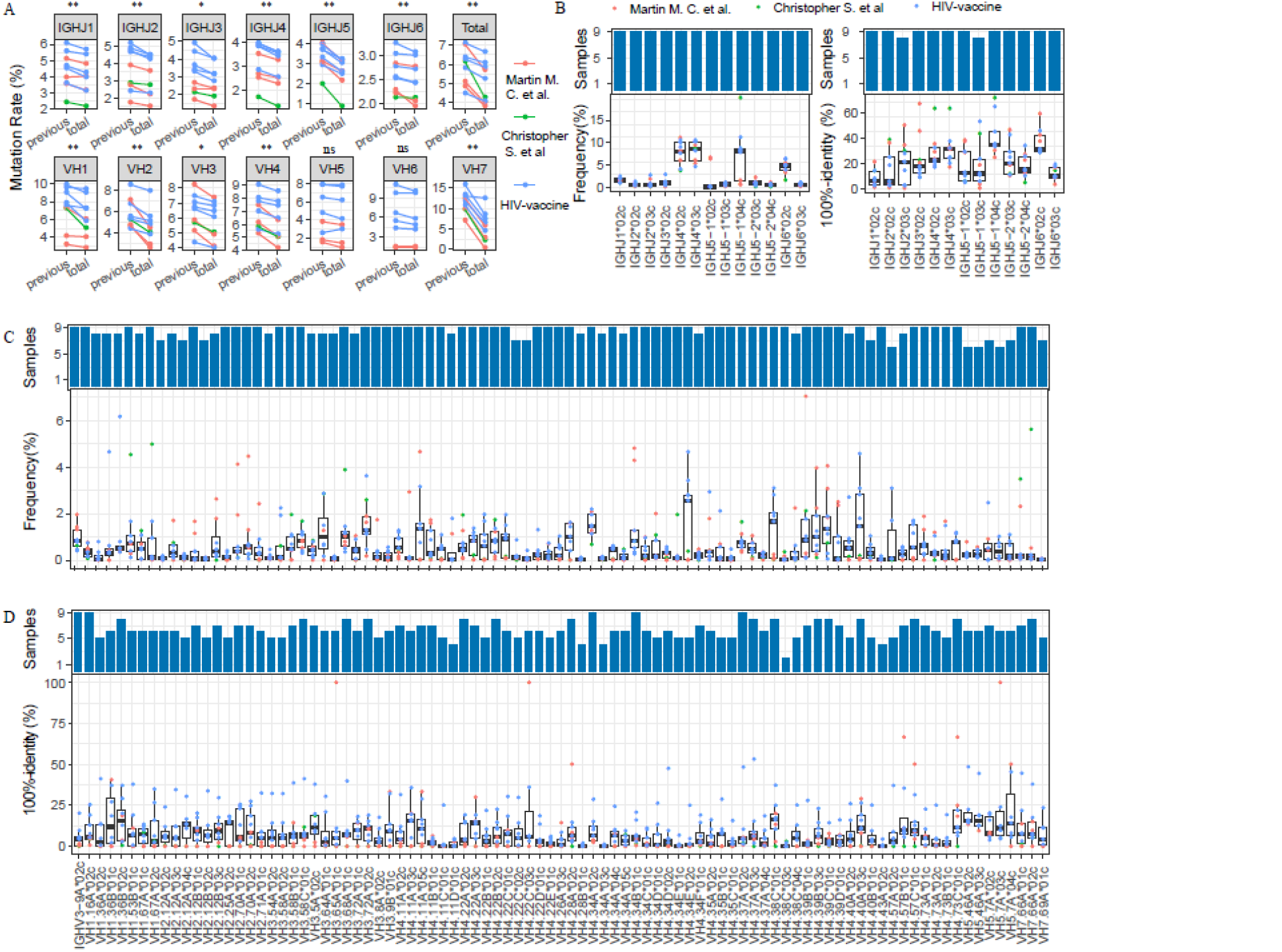
Novel germline alleles validated by other IgH repertoire data. (A). The average mutation rate for the VH and JH gene families. (*p<0.05, **p<0.01, ns, not-significant. paired Mann-Whitney test) (B). Sample number and mapped sequence frequency for each novel JH allele (left). The proportion of sequences (100% identity sequences mapped to the allele/total sequences mapped to the allele) mapped for each novel JH allele with 100% identity (right). (C). Sample number and mapped sequence frequency for each novel VH allele. (D). The proportion of sequences mapped for each novel VH allele with 100% identity.

Under the reference of MacVJ in repertoire analysis, we found that on average 53.59% and 31.33% of sequences could be mapped to the VH and JH novel alleles, respectively (Fig. S6B). Specifically, the sequence frequency that mapped to each novel allele are shown in Fig. 4B and Fig. 4D, respectively. All 9 samples contained sequences that were mapped to all 13 JH novel alleles, in which four alleles accounting for high frequencies (Fig. 4B). Although the frequency for each VH novel allele was present at a relatively low frequency, all 91 novel VH alleles were found in at least 6 samples (Fig. 4C). These results imply that rearranged sequences shared higher similarities with the novel VH and JH novel alleles than the previously reported alleles. However, to validate that the novel alleles were exactly correct, the mapping frequencies were inadequate. Therefore, we next looked into the mapped sequences that shared 100% identity with the novel alleles. We found that the average rates of 100%-identity sequences were 8.45% and 29.25% for the VH and JH, respectively (Fig. 6SB). We also displayed the rate and sample number for each novel JH and VH allele. All JH novel alleles were found in at least 8 samples that contained the 100% identity to the mapped sequences, and the average of 100%-identity rates for each JH allele ranged from 7.03% to 36.66% (Fig. 4B). Analogously, among the 100%-identity mapped sequences, all VH novel alleles were found in at least two samples, and 94.5% of alleles were observed in more than five samples, meanwhile, the average of 100%-identity rates for most (89%) VH alleles were more than 5% (Fig. 4D). Since every VH and JH novel alleles could be mapped with 100% identity, it strongly validates that the identification of novel alleles in our analysis is credible, lending further credence to the IMPre tool in repertoire sequence analysis.

### VH and JH gene Usage with MacVJ reference

To view the influence on the IgH repertoires analysis with our novel alleles, the sequencing data of 15 pre-vaccination Indian Rhesus samples were mapped to both reference datasets- the previously describe reference and MacVJ datasets. We utilized IMonitor to perform alignments and subsequent sequence analyses. In the previous references, most sequences were mapped to some predominated VH genes, such as VH4.11A, VH4.22A, VH4-34A, VH4.39B, VH4.40A (Fig. 5A). However, the VH gene usage in MacVJ displayed higher diversity, accompanied by the distribution of sequences on more genes. The frequencies of some genes had significant difference between two reference datasets. Moreover, some novel genes that were found in our study, such as VH4-39C, VH4-34B, VH4-11D, VH4-22B, VH4-22C, were characterized with relatively high frequencies (Fig. 5A). These results demonstrate that our inferred novel genes and alleles which mapped a large number of rearranged sequences have an important effect on the analysis of VH gene usage. As for the JH genes, although only novel alleles were added in comparisons with the MacVJ, some frequencies showed slight change in about 15 samples. These differences in frequencies in the JH genes included JH1, JH2, JH4, JH5-1 and JH5-2, respectively (Fig. 5B).

**Figure 5.**
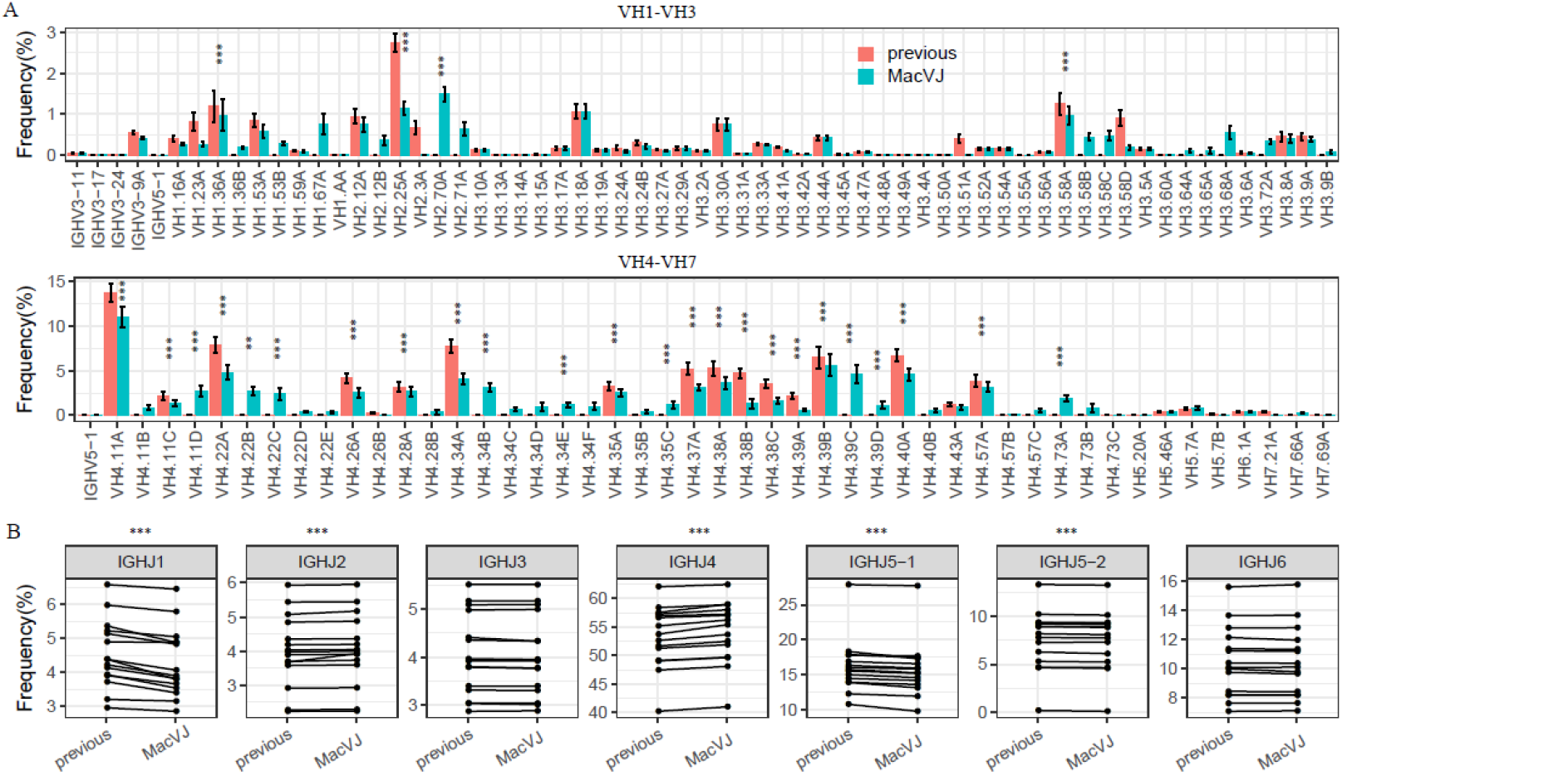
Rep-seq data of 15 India Rhesus were compared using previous and MacVJ datasets. (A). Bar plot for VH gene usage. Bars represent +/- standard error. Only the VH gene with average frequency of more than 1% was considered for the significance test. (B). JH gene usage. (**: p<0.01, p<0.001, paired Mann-Whitney test).

## Discussion

Advances in high-throughput sequencing technologies now allow for large-scale characterization of B cell immunoglobulin (Ig) repertoires, but the high germline and somatic diversity of the Ig repertoire poses challenges for biologically meaningful analysis, forcing the need for specialized computational methods. In this study, we utilized our developed software- IMpre and additional strategies to further process in multiple individuals to get the final novel VH and JH germline genes and alleles. We only selected the inferred germline genes and alleles that were present in multiple individuals in order to get a high-quality definition of the inferred alleles. The results from three human individuals analyzed herein demonstrated that this strategy coupled with the use of IMPre was reliable. Together, this method can be used by researchers to infer potential germline genes and alleles from large-scale rearranged data of B and T cell receptor repertoires. Using a series of aforementioned strict data filtration and evaluation strategies, we acquired 122 VH and 20 JH germline alleles, of which 91 VH alleles and 13 JH alleles were novel and have not been previously described. Two types of datasets were then used to validate these novel alleles. One dataset is the WGS reads of 10 Indian Rhesus monkeys, where we found fifty VH and five JH novel alleles. The other dataset is the Rep-seq reads which were from the previously published studies. We observed that all VH and JH novel alleles mapped to some rearranged sequences. Thus, these two independent datasets validated the high accuracy of inferred novel alleles observed and identified in our study. Besides, some Rhesus samples included in our analyses were treated with vaccine, but it has been well-documented that vaccine response varies, even in identical twins (27); therefore, vaccines should not affect germline gene/allele prediction.

We predicted more Rhesus VH4 alleles than other VH families. This could be attributed to the frequency of VH4, which was > 50% across samples in the Rep-seq data of Rhesus. Since there was a high proportion representing the VH4-containing sequences, other VH gene-containing sequences, which are relatively less frequent, were missed by the IMPre. In this context, it is important to iterate that principal utility of the IMPre software is based on the frequency of the gene segment. If the rearranged sequences derived from the VH germline allele accounting for very low frequency in the sample, this germline allele would not be inferred using the IMPre. Moreover, we cannot rule out the possibility that Rhesus monkeys may have more VH4 genes. One relevant finding is that the human germline VH3 contains more members than all other VH gene families taken together, and VH3-containing sequences account for the highest frequency among many rearranged sequence data sets (13, 27, 28). Likewise, VH4-containing sequences account for the highest percentage at approximately 50% in multiple monkey samples (2, 4). Since this estimation is partially speculative, more evidence is needed to validate this argument. We have also identified 8 novel genes located at a genome region with no previously reported Rhesus germline genes. Although some inferred alleles were mapped to a previously reported region, we also defined as novel genes when they contained more than a 7-bp mismatches. This threshold of 7 bp was derived from the analysis of human germline gene from a previously published study (21). We reiterate that this method may not be meticulous because of the incorrect classification of a portion of alleles into genes. Nonetheless, it is by far the best approach before a complete genome assembly from the Rhesus is available. In this study, we combined our inferred VH novel alleles, along with all previously reported VH germline alleles from the Indian Rhesus. However, the nomenclature of the genes and alleles had multiple versions, which needs a consensus from researchers working in this area. As for VH, we selected the rules of the KI_genome because they reported more genes. However, the nomenclature for the VH will become more accurate and informative for the VH positions in the Rhesus genome when the complete genome assembly is available.

it is a challenge to validate the novel VH and JH alleles, and there is no efficient and reliable method to do this. Even though primers can be designed in the novel allele and then be utilized to amplify the gene from non-lymphoid cells, it cannot ensure the amplification is exactly from germline VH or JH gene sets. The unique component of our work was using the two independent datasets to validate the novel alleles, which facilitated their accurate annotation. Most of the novel alleles are present in the Rhesus genome data, and some rearranged BCRs from other Rep-seq data are completely consistent with all novel alleles. Moreover, the mutation rate significantly declined after adding our novel alleles as the alignment reference. They could be observed in two other completely independent datasets, which imply that the novel alleles are correct and authentic. In addition, all our inferred novel alleles include a conservative amino acid motif at the start and end of the CDR3, which carries high similarity consistent with the previously reported Rhesus and Human germline alleles. Actually, these two aspects are the principal theories of finding KI_genome VH Rhesus germline alleles(19).

As more rearranged BCRs repertoires are sequenced in the future, the method inferring germline alleles from multiple individuals and proving novel alleles from WGS and Rep-seq data might prove useful in inferring more novel germline alleles. It is expected that with the completion of the Rhesus genome assembly in the near future, more VH and JH genes or alleles will be found and the nomenclature for the VH will become more precise. To add to this nomenclature precision will be the public availability of a comprehensive germline database created, which will significantly enhance the accuracy with which relationship between novel and already known germline alleles can be analyzed and inferred.

## Author Contributions

XL, WZ, IW designed the project; WZ developed the methodology and wrote the manuscript; LL, JW carried out the experiments; WZ designed the bioinformatic workflow; WZ, XY, LL, JX analyzed the data; IW, PQ, IM contributed reagents/materials; NK, XL, LT edited the manuscript. HM, JW provided funding support.

## Acknowledgments

This project was funded by the Beijing Genomics Institute and China National GeneBank. The funder provided support in the form of salaries for WZ, XY, LL, JX, LY, LT, JH and XL.

